# Gasdermin D mediates inflammation-induced defects in reverse cholesterol transport and promotes atherosclerosis

**DOI:** 10.1101/2021.04.16.440183

**Authors:** Emmanuel Opoku, C. Alicia Traughber, David Zhang, Amanda J Iacano, Mariam Khan, Juying Han, Jonathan D Smith, Kailash Gulshan

## Abstract

Activation of inflammasomes, such as Nlrp3 and Aim2, can exacerbate atherosclerosis in mice and humans. Gasdermin D (GsdmD) serves as a final executor of inflammasome activity, by generating membrane pores for the release of mature Interleukin-1beta (IL-1β). Inflammation dampens reverse cholesterol transport (RCT) and promotes atherogenesis, while anti-IL-1β antibodies were shown to reduce cardiovascular disease in humans. Though Nlrp3/AIM2 and IL-1β nexus is an emerging atherogenic pathway, the direct role of GsdmD in atherosclerosis is not yet fully clear. Here, we used in-vivo Nlrp3 inflammasome activation to show that the GsdmD^-/-^ mice release ~80% less IL-1β vs WT mice. The GsdmD^-/-^ macrophages were more resistant to Nlrp3 inflammasome mediated reduction in cholesterol efflux, showing ~26% decrease vs. ~60% reduction in WT macrophages. GsdmD expression in macrophages exacerbated foam cell formation in an IL-1β dependent fashion. The GsdmD^-/-^ mice were resistant to Nlrp3 inflammasome mediated defect in RCT, with ~32% reduction in plasma RCT vs. ~ 57% reduction in WT mice, ~ 17% reduction in RCT to liver vs. 42% in WT mice, and ~ 37% decrease in RCT to feces vs. ~ 61% in WT mice. The LDLr anti-sense oligonucleotides (ASO) induced hyperlipidemic mouse model showed the role of GsdmD in promoting atherosclerosis. The GsdmD^-/-^ mice exhibit ~42% decreased atherosclerotic lesion area in females and ~33% decreased lesion area in males vs. WT mice. The atherosclerotic plaque-bearing sections stained positive for the cleaved N-terminal fragment of GsdmD, indicating cleavage of GsdmD in atherosclerotic plaques. Our data show that GsdmD mediates inflammation-induced defects in RCT and promotes atherosclerosis.

**Summary:** GsdmD mediates inflammation-induced defects in RCT and promotes atherosclerosis.

## Introduction

The NLRP3 inflammasome is implicated in promoting cardiovascular disease (CVD) and metabolic diseases such as obesity-induced inflammation/insulin resistance (*1, 2*), and destabilization of atherosclerotic plaques (*1, 3*). The Canakinumab Anti-Inflammatory Thrombosis Outcomes Study (CANTOS) trial showed that anti-IL-1β therapy reduced major adverse coronary events (MACE), independent of lipid levels (*4*). The Nlrp3 inflammasome target protein “GsdmD” is involved in multiple pro-inflammatory pathways such as the release of IL-1β, pyroptotic cell death, and formation of neutrophil extracellular traps/NETosis (*5, 6*). Nlrp3 inflammasome is activated in advanced human atherosclerotic plaques, but the role of GsdmD in sterile inflammatory diseases such as atherosclerosis is not yet described. Interestingly, GsdmD cleavage does not necessarily lead to pyroptotic cell death under all conditions, as GsdmD also plays a role in the release of IL-1β from living macrophages (*7–9*). Thus living, but inflamed, macrophages in atherosclerotic plaques may contribute to elevating IL-1β levels in GsdmD dependent manner.

Deposition and oxidative modification of low-density lipoprotein-cholesterol (LDL-C) in the arterial intima promotes monocyte entry and transformation into macrophages, leading to lipid engulfment and foam cell formation. Foam cells become dysfunctional over time due to unregulated lipid uptake, leading to fatty streaks and further amplification of inflammation and progression of atherosclerosis (*1, 10–12*). Removal of excess cholesterol from artery wall macrophages via reverse cholesterol transport (RCT) may reverse the progression of atherosclerosis (*13, 14*). Chronic inflammation serves as a double-edged sword by promoting the continued influx of immune cells into the plaque area via inducing the expression of adhesion molecules such as vascular cell adhesion molecule-1 (VCAM-1) on endothelial cells, and by dampening the protective RCT pathway (*15, 16*). Accumulation of oxidized lipids and cholesterol crystals in plaques can engage the toll-like receptor (TLR) pathway and induce the assembly of the NLRP3 inflammasome (*1, 3, 12, 17*). The NLRP3 inflammasome plays a key role in processing procaspase 1 to active caspase 1, which in turn can mediate the cleavage of pro-interleukin-1β (pro-IL-1β), pro-interleukin-18 (pro-IL-18), and Gasdermin D (GsdmD) to generate active IL-1β, active IL-18, and active N-terminal fragment of GsdmD (GsdmD-NT). Mature IL-1β can be released from inflamed macrophages in at least two ways. In one pathway, the cleaved GsdmD-NT binds to phosphatidylinositol lipids (PIPs) and phosphatidylserine (PS) on the inner leaflet of the plasma membrane of cells to generate pores for the fast release of mature IL-1β (*18–20*). In the second pathway, the cleaved IL-1β exit via directly binding to the phosphatidylinositol 4,5-bisphosphate (PIP2) in plasma membrane lipid rafts (*21*). In contrast to Nlrp3 inflammasome and IL-1β, the direct role of GsdmD in atherosclerosis is not yet clear. Though GsdmD cleavage and ensuing membrane blebbing and pyroptosis are generally associated with the robust release of cytokines and clearance of microbial infection, it’s important to note that GsdmD also plays a role in IL-1β release from living macrophages (*22*). Furthermore, similar to the apoptotic death of cholesterol-loaded macrophages (*23, 24*), the pyroptotic cell death can be induced by cholesterol crystals mediated Nlrp3 inflammasome activation in advanced human atherosclerotic plaques (*1, 25*). Thus, GsdmD may play role in Nlrp3 inflammasome mediated dampening of RCT and in promoting the progression of atherosclerosis, either by virtue of increasing IL-1β release from live inflamed immune cells or by promoting pyroptotic cell death in advanced atherosclerotic plaques.

## Results

### GsdmD^-/-^ mice release lower levels of IL-1β upon in-vivo Nlrp3 inflammasome assembly

To confirm that GsdmD ^-/-^ macrophages are defective in IL-1β release, primary bone-marrow-derived macrophages (BMDMs) isolated from wild type (WT) or GsdmD^-/-^ mice were primed with 1μg/ml LPS for 4h. To induce Nlrp3 inflammasome assembly, these cells were incubated with 5mM ATP for 20 min or 1 μM Nigericin for 1h, followed by a collection of cell-free media to measure released IL-1β. Consistent with previous studies, the GsdmD^-/-^ macrophages showed a ~80% decrease in the release of IL-1β as compared to control macrophages (***Fig. 1A***). The basal as well as the LPS induced expression levels of NLRP3 and pro-IL-1β were not altered in GsdmD^-/-^ BMDMs, indicating that only IL-1β release is defective (***Fig 1B***).

**Fig. 1.**
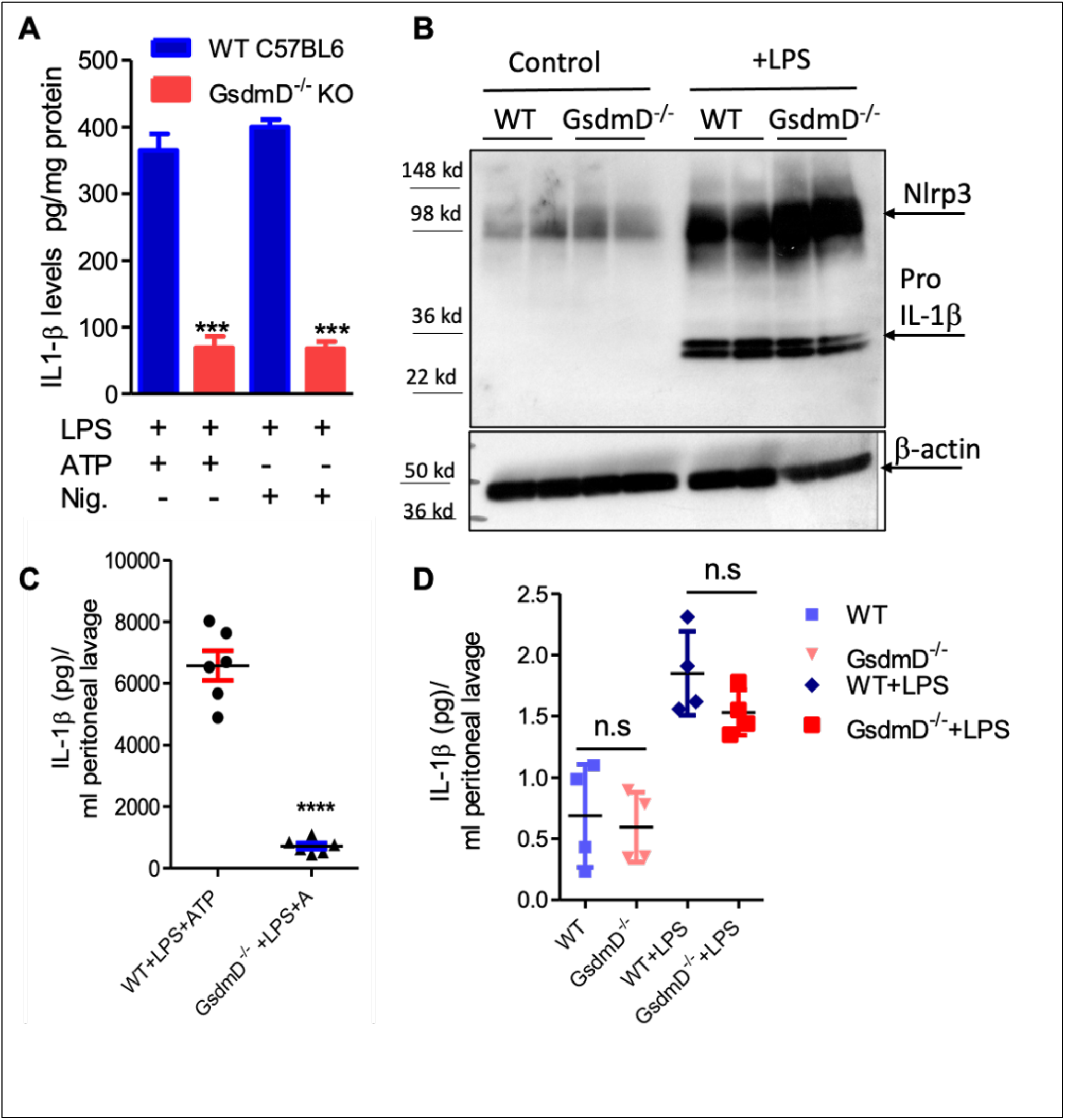
*A)* GsdmD plays a role in release of mature IL-1β upon in-vivo Nlrp3 inflammasome activation. ***A***) BMDMs isolated from WTC57BL/6J or C57BL6/J-GsdmD^-/-^ mice were primed by incubation with ± 1μg/ml LPS for 4h. The Nlrp3 inflammasome assembly was induced either by incubation with 1 mM ATP for 20 min or 1μM Nigericin for 1h. The IL-1β levels in the cell-free media were measured by ELISA (mean ± SD, N = 6, *** represent p<0.001 by ANOVA with Bonferroni posttest). ***B***) WT or GsdmD^-/-^ BMDMs were treated ± 1 μg/ml LPS, followed by western blot analysis using Nlrp3 and-IL-1β antibodies. The β-actin was used as a loading control. ***C***) The age-matched male WTC57BL/6J or GsdmD^-/-^ mice were primed with IP injection of LPS (5μg/mouse). After 4h of LPS injection, the Nlrp3 inflammasome assembly in mice was induced by IP injection of ATP (0.5 ml of 30 mM, pH 7.0). The peritoneal cavity was lavaged with 3 ml PBS, and IL-1β levels in peritoneal lavage were determined by ELISA (N=6, **** p <0.0001 by two-tailed t-test). **D**) The age-matched male WTC57BL/6J or GsdmD^-/-^ mice were treated with ± IP injection of LPS (5μg/mouse) and injected with 0.5 ml of saline. The peritoneal cavity was lavaged with 5 ml PBS, and IL-1β levels in peritoneal lavage were determined by ELISA (N=4, n.s.=non-significant by two-tailed t-test).

To determine the role of GsdmD in Nlrp3 inflammasome mediated IL-1β release in-vivo, the Nlrp3 inflammasome was induced in mice by i.p. injection of 5 μg LPS, followed 4h later with i.p injection of 0.5 ml of 30 mM ATP. Control mice received either saline or LPS + saline injections. The peritoneal lavage fluid was collected 30 min after ATP injection and analyzed for IL-1β levels by ELISA. As shown in ***Fig. 1C***, WT mice injected with LPS and ATP showed robust IL-1β levels in peritoneal lavage while the GsdmD^-/-^ mice had a ~ 90 % reduction in IL-1β levels. The IL-1β levels were not detectable in peritoneal lavage of WT or GsdmD^-/-^ mice injected with saline or LPS alone (***Fig. 1D***), showing that both LPS and ATP are required for *in vivo* Nlrp3 inflammasome activation and GsdmD is required for IL-1β release.

### GsdmD^-/-^ macrophages showed protection from Nlrp3 inflammasome-induced defect in cholesterol efflux

To determine if Nlrp3 inflammasome activation leads to defective cholesterol efflux and if GsdmD deletion protects against this effect, cholesterol efflux assays were performed in BMDMs ± induction of Nlrp3 inflammasome. The BMDMs were labeled with ^3^H cholesterol for 24 h, followed by induction of ABCA1 expression with liver X receptor (LXR) agonist T-0901317. The macrophages were treated ± LPS and Nigericin, followed by a chase with 5μg/ml lipid-free apoA1 for 4h. As shown in ***Fig. 2A***, LPS/Nigericin treatment of WT BMDMS led to ~ 60% decrease in cholesterol efflux vs. untreated BMDMs. The BMDMs derived from GsdmD^-/-^ KO mice showed much lower inhibition of cholesterol efflux (~26% decrease vs. untreated). The direct effect of IL-1β was evaluated by treatment of LPS treated WT or GsdmD^-/-^ BMDMs with exogenous recombinant mouse IL-1β. The addition of exogenous recombinant mouse IL-1β inhibited cholesterol efflux in both WT and GsdmD^-/-^ BMDM (***Fig. S1***). These data indicated that exogenous IL-1β is sufficient to drive GsdmD^-/-^ KO macrophages to respond similarly to WT macrophages. Thus, IL-1β appears to be a major factor responsible for inhibiting cellular cholesterol efflux. To recapitulate the environment of atherosclerotic plaque area, where IL-1β released from inflamed foam cells may affect the cholesterol efflux capacity of neighboring cells, we used the conditioned media from BMDMs treated with LPS/ATP. The conditioned media from WT BMDMs inhibited cholesterol efflux in both WT and GsdmD^-/-^ KO macrophages, while the conditioned media from GsdmD^-/-^ BMDMs treated with LPS/ATP showed reduced suppression of cholesterol efflux (***Fig. 2B, S1***). These data indicated that macrophages lacking GsdmD are more resistant to inflammasome-induced reduction in cholesterol efflux, mostly due to defective IL-1β release. The ABCA1 expression was pre-induced with T-compound, and as expected no significant differences in ABCA1 levels were observed in control or conditioned-media treated macrophages (***Fig. S2***). Next, we tested if IL-1β is interfering with apoA1 binding to the cell membrane. The apoA1 binding to the cellsurface ± WT conditioned media was probed by using an Alexa647 labeled apoA1 as described before (*26, 27*). The BMDMs showed basal apoA1 binding as they express ABCA1, albeit at a low level, while induction of ABCA1 with T compound led to a significant increase in apoA1 binding to the cell surface (***Fig. 2C. 2D***). Treatment of BMDMs with WT-CM led to significant decrease in apoA1 binding to the cell surface (***Fig. 2C, 2D***), while the GsdmD^-/-^ CM media showed no effect on apoA1 binding to the cell surface (***Fig. 2D)**.*

**Fig. 2.**
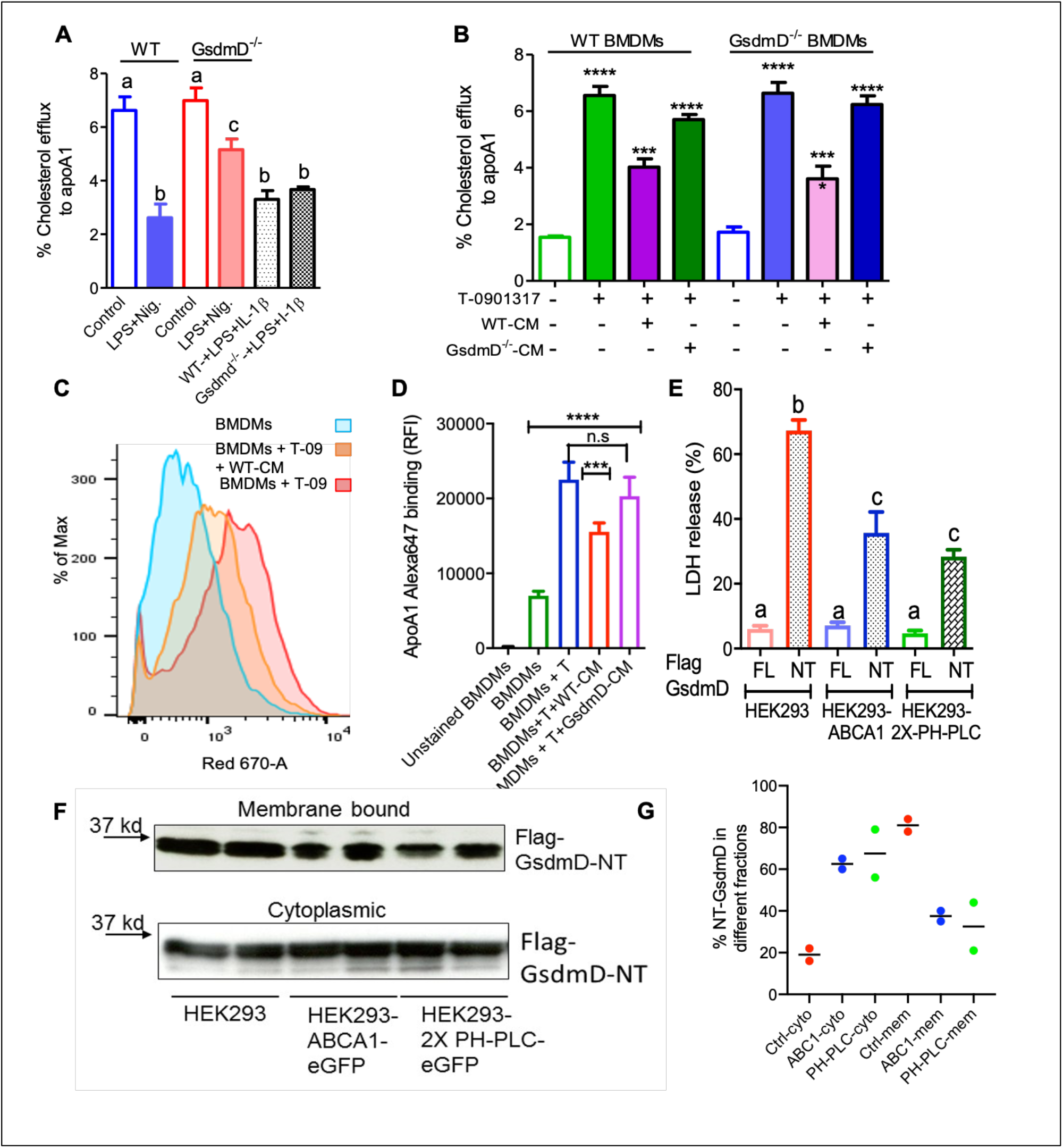
GsdmD promotes Nlrp3 inflammasome-mediated defects in reverse cholesterol transport. ***A***) BMDMs isolated from WT or GsdmD^-/-^ mice were loaded with ^3^H-cholesterol + 50μg/ml AcLDL for 24h and treated with LXR-agonist T0901317 (Sigma-Aldrich) to induce ABCA1 expression. The Nlrp3 inflammasome was induced by LPS+ Nigericin treatment, followed by cholesterol efflux assay using lipid-free apoA1 (5μg/ml) in serum-free DMEM as chase media for 4h at 37°C. Values are % cholesterol efflux (mean ± SD, N = 4, **** represent p<0.0001, ** represent p<0.01 by ANOVA Bonferroni posttest). ***B***) The BMDMs isolated from WT or GsdmD^-/-^ mice were loaded with ^3^H-cholesterol + 50μg/ml AcLDL for 24h. The BMDMs, ± LXR-agonist T0901317, were treated with conditioned media isolated from LPS+ATP treated WT or GsdmD^-/-^ BMDMs for 16h. The cholesterol efflux assay was performed using lipid-free apoA1 (5μg/ml) in serum-free DMEM as chase media for 4h at 37°C. Values are % cholesterol efflux (mean ± SD, N = 4, **** represent p<0.0001, *** represent p<0.001 by ANOVA Bonferroni posttest). ***C***) The flow cytometry analysis showing histogram bar for Alexa-647 labeled apoA1 binding to cells, ± ABCA1 expression and ± WT conditioned media treatment ***D***) The flow cytometry analysis showing quantification of Alexa-647 labeled apoA1 binding to cells ± ABCA1 expression and ±. WT or GsdmD^-/-^ conditioned media treatment (mean ± SD, N = 4, **** represent p<0.0001, *** represent p<0.001 by ANOVA Bonferroni posttest) ***E***) The HEK293, HEK293-ABCA1-eGFP, or HEK293-2X-PH-PLC-eGFP cells were transfected with plasmids carrying either full-length (FL) GsdmD or N-terminal fragment (NT) of GsdmD. The LDH release in media is plotted in % values of total cellular LDH (mean ± SD, N = 4; different letters show p<0.001, by ANOVA Bonferroni posttest). ***F***) The control HEK293 and HEK293 cells stably transfected with ABCA1 or PH-PLC were transfected with plasmid carrying Flag tagged GsdmD-NT. The transfected cells were fractionated into cytoplasmic or membrane bound fractions. The fractions were resolved using SDS gel, transferred to nitrocellulose membrane, and probed with antibody against Flag tag.

### PIP2 localization modulated GsdmD-NT mediated cell lysis

PIP2 serves as a ligand for GsdmD N-terminal (GsdmD-NT) fragment and transfection of GsdmD-NT in HEK293 cells promotes cell lysis (*18*). ABCA1 expression reduces PIP2 on the inner leaflet of plasma membrane (*27, 28*), thus we tested if ABCA1 expression can reduce GsdmD mediated cell lysis by transfecting the control HEK293 and HEK293-ABCA1 cells with GsdmD-NT. Transfection of GsdmD-NT, but not the full-length GsdmD, caused cell lysis in HEK293 cells (***Fig. 2E***). The HEK293 cells expressing ABCA1 showed reduced cell lysis (***Fig. 2E***), indicating that PIP2 flop across the cell membrane can dampen GsdmD-NT mediated pore-formation and cell lysis. To test the direct role of PIP2 in GsdmD-NT cell lysis, we generated HEK293 cells stably expressing GFP tagged Pleckstrin homology domain of Phospholipase-C protein (2X-PH-PLC-eGFP). As the majority of cellular PIP2 is localized to the inner leaflet of plasma membrane, as expected the PH-PLC-eGFP signal was localized at the plasma membrane (***Fig. S3***). The HEK293-2X-PH-PLC-eGFP cells transfected with GsdmD-NT showed reduced lysis vs. control HEK293 cells (***Fig. 2E).*** The membrane recruitment of N-terminal GsdmD was confirmed by fractionation of cytosolic and membrane proteins using the sequential fractionation method as described earlier (*29*). The membrane recruitment of GsdmD-NT was reduced in HEK-ABCA1 cells and HEK-PH-PLC cells vs. control HEK293 cells (***Fig. 2F, 2G***). The control HEK293 cells showed ~80 % of GsdmD-NT recruitment to the membrane while the rest ~ 18% remained in cytoplasm. In contrast, the HEK-ABCA1 and HEK-PH-PLC cells showed ~ 40 % and ~ 35% recruitment of GsdmD-NT to the membrane, while ~ 60% and ~ 65% remains in cytoplasm. These data indicated that blocking PIP2 access to NT-GsdmD reduced recruitment of GsdmD-NT to the plasma membrane, leading to lower cell lysis.

### GsdmD promotes foam cell formation

To determine if the IL-1β released from inflamed cells can promote foam cell formation, the lipid loading was performed along with treatment with conditioned media. The WT BMDMs were incubated with the conditioned media isolated from LPS+ATP treated WT or GsdmD^-/-^ macrophages along with 25 μg/ml Acetylated-low-density lipoprotein (AcLDL) and 25 μg/ml oxidized low-density lipoprotein (oxLDL) for 48h to induce foam cell formation. The BMDMs were washed thoroughly with serum-free media and stained with lipid-binding Nile-red dye, followed by either qualitative analysis via fluorescent microscopy or quantitative analysis via flow cytometry. The WT BMDMs, as well as the GsdmD^-/-^ BMDMs, showed higher lipid accumulation in presence of WT conditioned media (***Fig. 3A, Fig. S5***). In contrast, the WT or GsdmD^-/-^ BMDMs showed reduced foam cell formation with GsdmD^-/-^ conditioned media treatment (***Fig. 3A, 3B)**.* These data indicated that GsdmD mediated IL-1β release promotes foam cell formation. Since the GsdmD^-/-^ macrophages are resistant to pyroptotic cell death, we sought to determine if lipid-laden GsdmD^-/-^ can instead undergo apoptotic cell death. Foam cell formation was induced by loading WT or GsdmD^-/-^ BMDMs with 100 μg/ml AcLDL for 48h, followed by Nlrp3 inflammasome activation by LPS and ATP treatment. The cells were washed with PBS and the phosphatidylserine (PS) exposure at the cell surface was determined by flow cytometry based Annexin-Cy5 binding assay as described earlier(*27*). As shown in ***Fig. 3C***, the GsdmD^-/-^ foam cells showed a much higher annexin V signal vs. WT foam cells (N=4, mean ± SD). These data indicated that the lack of GsdmD in Nlrp3 inflammasome-induced macrophages can promote apoptotic cell death in lieu of pyroptosis.

**Fig. 3.**
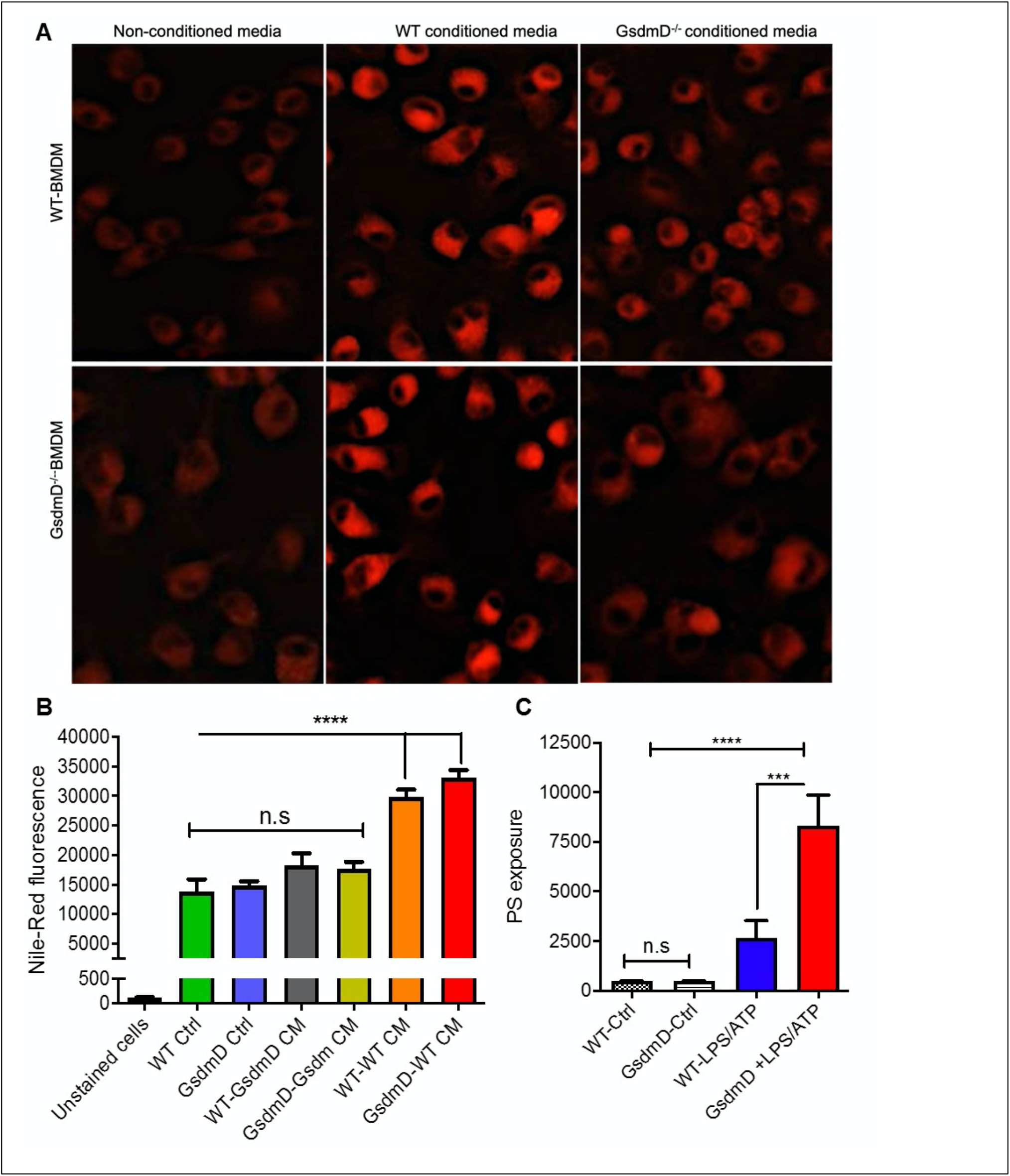
GsdmD promotes foam cell formation. ***A***) BMDMs isolated from WT or GsdmD^-/-^ mice were loaded with 50μg/ml AcLDL + 25μg/ml OxLDL for 48 h ± conditioned media isolated from LPS/ATP treated WT or GsdmD^-/-^ BMDMs. The culture media was removed, cells were washed with PBS, and stained with Nile-red dye (0.25 μg/ml) at 37°C for 15 min. The stained cells were washed thoroughly with PBS and fixed with paraformaldehyde and imaged using fluorescent microscope with TRITC filter (Excitation / emission: 550 / 640 nm). ***B***) Quantification of Nile-red staining in WT or GsdmD^-/-^ BMDMs ± conditioned media by flow-cytometry, mean ± SD, N = 4, **** represent p<0.0001 by ANOVA Bonferroni posttest. ***C***) BMDMs isolated from WT or GsdmD^-/-^ mice were loaded with 50μg/ml AcLDL + 25μg/ml OxLDL for 48 h. The foam cells were left untreated or treated with LPS+ATP to induce Nlrp3 inflammasome. The cells were washed with PBS, and PS exposure was determined by flow-cytometry based assay using Annexin Cy5 staining (mean ± SD, N = 4, **** represent p<0.0001, *** represent p<0.001 by ANOVA Bonferroni posttest, n.s=non-significant).

### GsdmD^-/-^ mice showed protection from Nlrp3 inflammasome-induced defect in RCT

To determine if Nlrp3 inflammasome assembly in mice leads to defective RCT and if GsdmD^-/-^ KO mice are protected against this effect, an RCT assay was performed as described earlier (*16*). The BMDMs from WT mice were loaded with acetylated-low-density lipoprotein (Ac-LDL) and radiolabeled ^3^H cholesterol for 48 h to generate foam cells. The WT and GsdmD KO mice pretreated with LPS + ATP treatment were injected with foam cells, following the RCT assay design shown in ***Fig. 4A***. The radioactive cholesterol efflux to plasma was determined by collection of blood samples at 24h, 48h, and 72h. The liver and feces samples were collected post euthanasia and processed to determine the % of cholesterol efflux. As shown in ***Fig. 4B***, Nlrp3 inflammasome activation reduces RCT to plasma in WT mice by 67% at 24 h, by 33% at 48 h, by 47% at 72 h (Fig. 2B), while GsdmD^-/-^ mice showed only 46% reduction at 24 h, 26% at 48 h, and 33% reduction at 72 h (***Fig. 4C***). The cumulative reduction in RCT to plasma in WT mice was ~57% vs. 32% in GsdmD^-/-^ mice, showing that the GsdmD^-/-^ mice have 25% more preserved RCT to plasma vs. WT mice (***Fig. 4D***). The RCT to liver was reduced by 42% in WT mice vs. 17% reduction in GsdmD^-/-^ mice (***Fig. 4E***). The RCT to feces was reduced upon Nlrp3 inflammasome activation by 31% at 24 h, by 77% at 48 h, by 63% at 72 h in WT mice (***Fig. 4F***), while GsdmD^-/-^ mice showed 18% reduction at 24 h, 40% at 48 h, and 38% reduction at 72 h (***Fig. 4G***). The cumulative reduction in RCT to feces in WT mice was ~61% vs. 37% in GsdmD^-/-^ mice, showing that GsdmD^-/-^ mice have 24% more preserved RCT to feces vs. WT mice upon Nlrp3 inflammasome induction (***Fig. 4H***). These data indicated that GsdmD promotes inflammation-induced defects in reverse cholesterol transport.

**Fig. 4:**
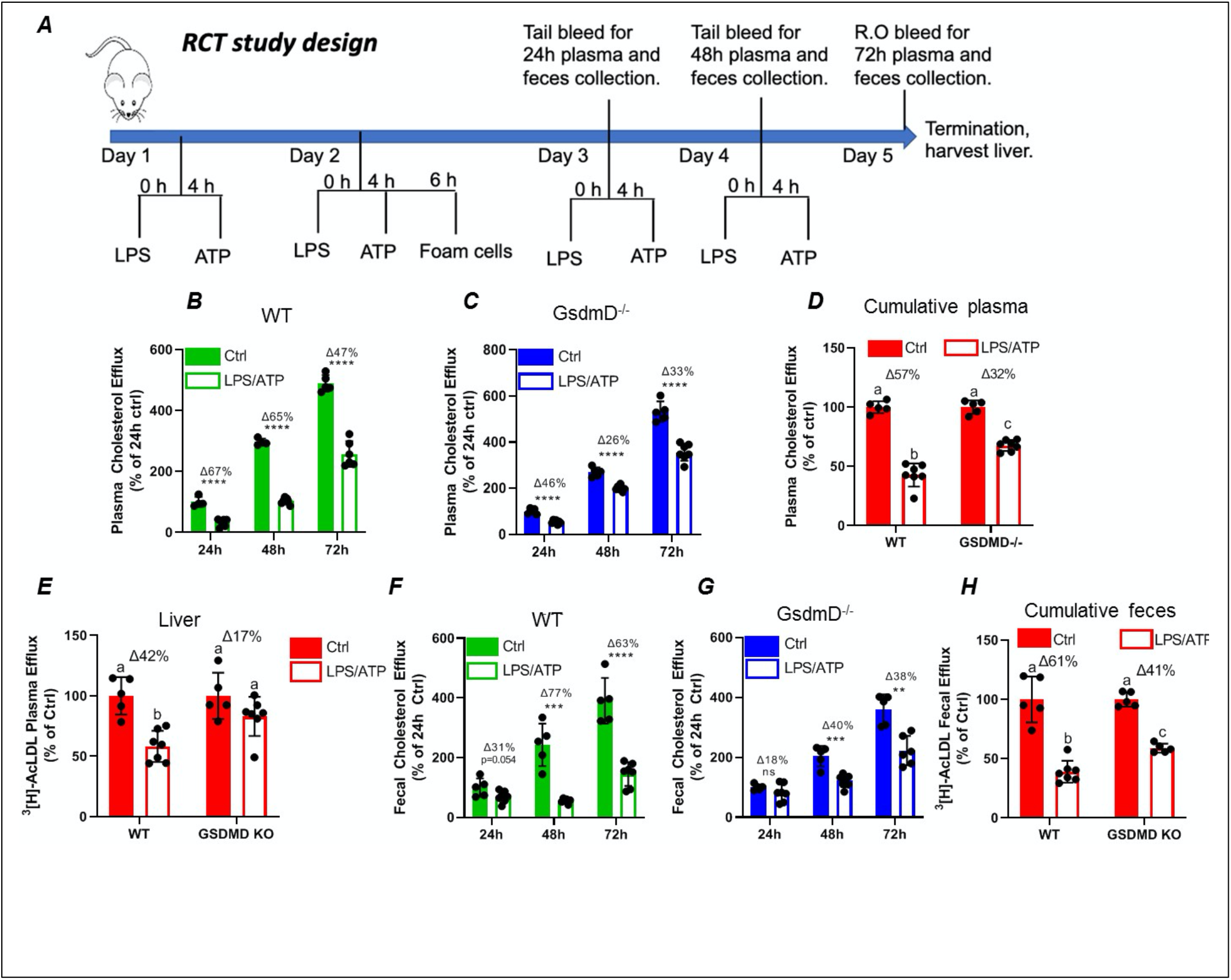
GsdmD mediates Nlrp3 inflammasome-induced defects in reverse cholesterol transport. ***A***) Schematic representation of mouse RCT study design. The 12-week-old male mice were injected daily with or without 5ug LPS and 30mM ATP for 4 days. On day 2 mice were injected with radiolabeled foam cells loaded with AcLDL and ^3^H cholesterol. The plasma and feces RCT was assessed at 24,48,72h (quantified as a % of the 24h Ctrl for each group) and cumulatively (quantified as a percent of respective Ctrl). ***B***) RCT to plasma in WT mice at 24h, 48h, and 72h. ***C***) RCT to plasma in GsdmD^-/-^ mice at 24h, 48h, and 72h. ***D***) Cumulative RCT to plasma in WT and GsdmD^-/-^ mice. ***E***) RCT to liver in WT and GsdmD^-/-^ mice. ***F***) RCT to feces in WT mice at 24h, 48h, and 72h. ***G***) RCT to feces in GsdmD^-/-^ mice at 24h, 48h, and 72h. ***H***) Cumulative RCT to feces in WT and GsdmD^-/-^ mice. N=4-5, for WT and GsdmD^-/-^ Ctrl mice, N=6-7 for WT and GsdmD^-/-^ LPS/ATP mice. ****p<0.0001, ***p<0.001, **p<0.01 by two-tailed t-test. Groups with different letters a, b, c, d is p<0.05 by One-way ANOVA.

### LDLr ASO induces hyperlipidemia in WT and GsdmD^-/-^ mice

Antisense oligonucleotides directed to LDLR mRNA cause hypercholesterolemia in western type diet (WTD) fed wild-type C57BL/6 mice (*30*). The 12-week-old WT and GsdmD^-/-^ KO mice were injected i.p with LDLr ASO or control ASO once a week for 9 weeks and mice were fed an atherogenic WTD throughout the study, as shown in the atherosclerosis study design (***Fig. 5A***). The mice were weighed and plasma was collected before and after the 9-week ASO treatments, and total cholesterol (TC) levels were determined. As shown in ***Fig. 5B,*** no significant differences were found in body weight or cholesterol levels in WT and GsdmD^-/-^ female mice before or after the ASO and diet interventions. The plasma TC levels showed a robust increase in LDLr ASO treated female mice and reached an average level of ~ 800 mg/dl (Fig. ***5C***). The male mice also showed similar trends with no difference in body weight gain and increase in plasma cholesterol upon LDLr ASO treatment (***Fig. 5D, 5E***). These data showed that the LDLr ASO has equivalent efficacy in both WT and GsdmD^-/-^ mice.

**Fig. 5.**
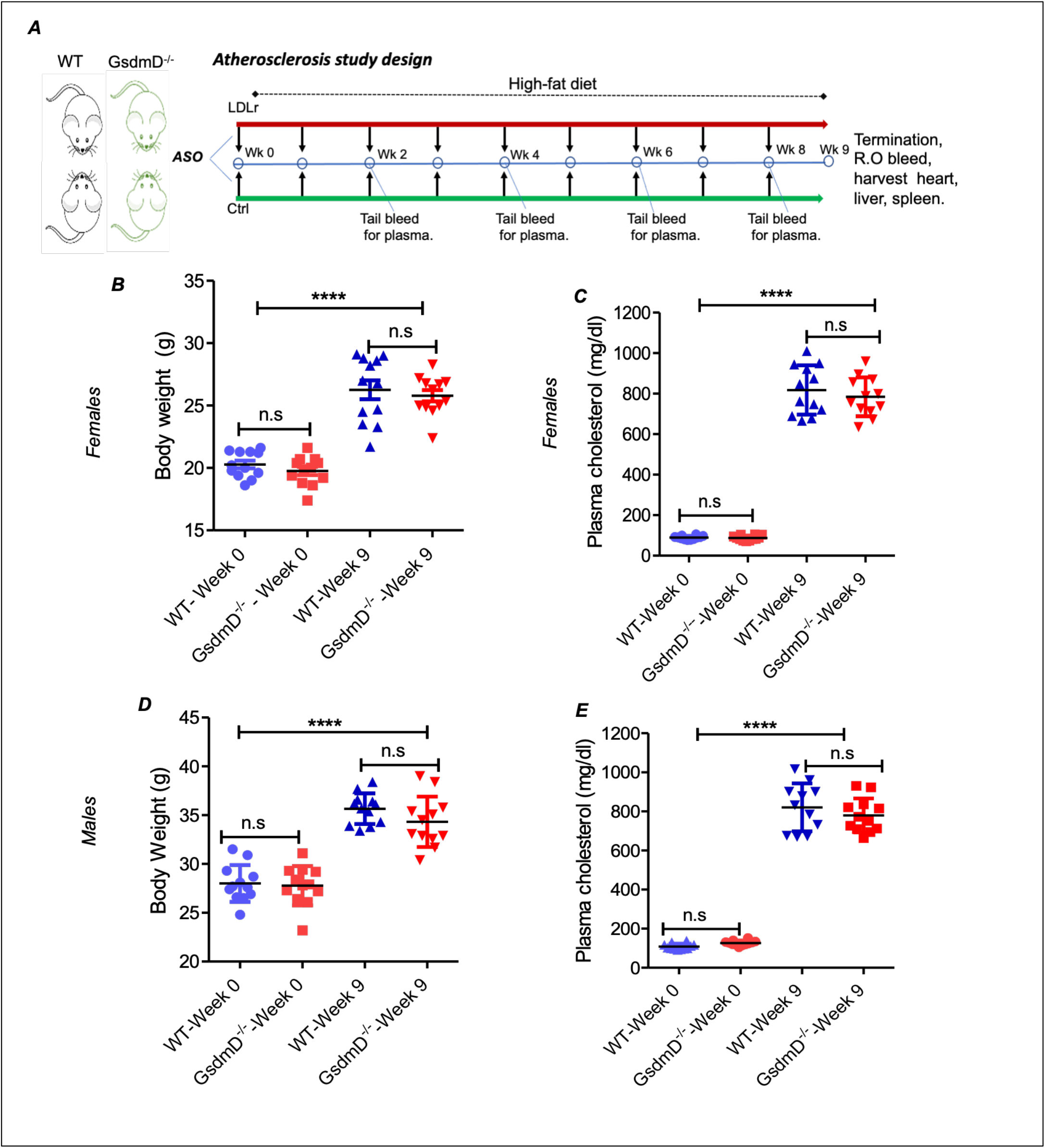
LDLr ASO generates hyperlipidemia in both WT and GsdmD^-/-^ KO mice. ***A***) Schematic representation of mouse atherosclerosis study design. The 12-week-old mice from both sexes were injected with either control or LDLr ASO for 9 weeks. The mice were fed western type diet throughout the course of study. The body weight in females (***B***) and total plasma cholesterol levels (***C***) at week 0 and at the end of experiment, indicated by week 9, (mean ± SD, N = 12, **** p <0.0001 by two-tailed t-test, n.s= non-significant), and male data for body weight (**D**) and total plasma cholesterol levels (***E***) is presented at base and final levels (N=12 **** p <0.0001 by two-tailed t-test for both WT-week 0 vs. WT-week 9 and GsdmD^-/-^-week 0 vs. GsdmD^-/-^-week 9, n.s= non-significant by two-tailed t-test).

### GsdmD^-/-^ mice showed reduced atherosclerosis

LDLr ASO treatment and WTD feeding lead to the development of atherosclerotic lesions in the aortic root, aortic arch, and brachiocephalic artery (*30*). To determine if GsdmD plays a role in the progression of atherosclerosis, the 12-week-old WT and GsdmD^-/-^ mice were injected i.p with LDLr ASO or control ASO once a week, and fed with atherogenic western-type diet (WTD), to generate hyperlipidemia. The mice were sacrificed at the end of week 21 and hearts were perfused with PBS, followed by harvesting of heart, spleen, and liver. The knockdown of hepatic LDLr by the LDLr ASO was confirmed by the western blot of liver extracts (***Fig. 6A***).

**Fig. 6.**
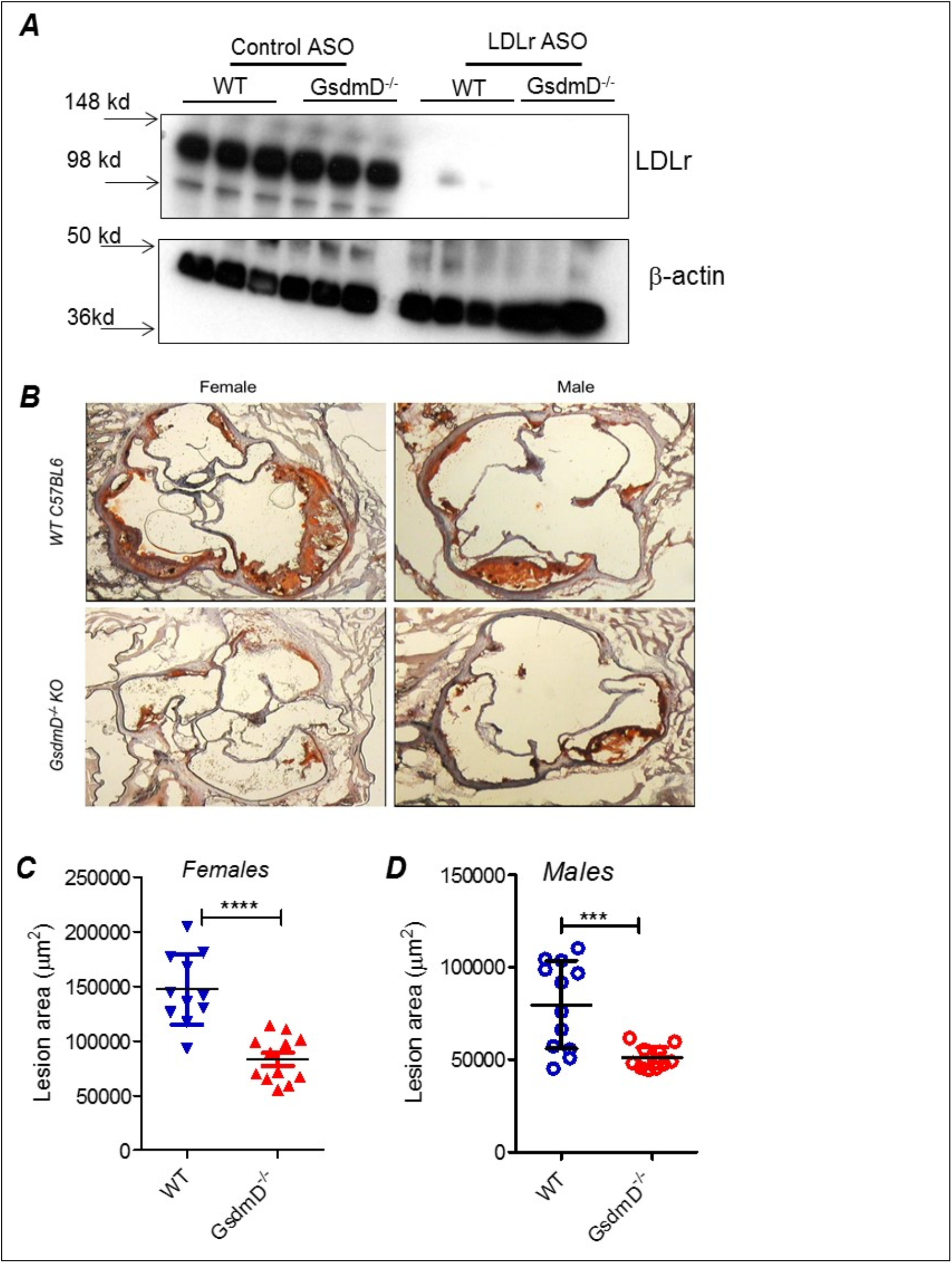
GsdmD promotes atherosclerosis and is cleaved in atherosclerotic plaques. The 12-weeks old mice from both sexes were injected with either control or LDLr ASO for 9 weeks and were fed western type diet throughout the course of study. Mice were euthanized after 9-week ASO treatment and were perfused with PBS and hearts and livers were removed. ***A)*** A small portion of liver from mice was excised, weighed, and protein extracts were prepared using RIPA buffer supplemented with protease inhibitors. Western blot analysis of LDLr in liver extracts was performed using LDLr antibody and HRP conjugated secondary antibody. ***B)*** Representative images from both sexes showing aortic root lesions stained with Oil red O and hematoxylin from WT or GsdmD^-/-^ KO mice injected with LDLr ASO and fed WTD, ***C, D)*** Quantification of aortic root lesions in females and males from WT or GsdmD^-/-^ KO mice treated with LDLr ASO and fed WTD (for WT male N=11, for WT females, GsdmD^-/-^ males, and GsdmD^-/-^females, N=12, *** p <0.0005, **** p<0.0001 by two-tailed t-test).***E)*** Protein extracts from aortas of control ASO or LDLr ASO treated mice (N=4) were subjected to western blot analysis using antibody specific for cleaved GsdmD and VCAM-1. GAPDH was probed as a loading control. ***F)*** ATP levels in plasma of control ASO or LDLr ASO treated mice fed with atherogenic diet were determined by ATP quantification kit, following manufacturer’s instructions. (mean ± SD, N = 8; different letters show p<0.001, by ANOVA Bonferroni posttest).

The hearts were sectioned and the atherosclerotic plaques were quantified using the oil red O positive aortic root atherosclerotic lesion area. As shown in ***Fig. 6B***, WT mice injected with LDLr ASO showed large atherosclerotic lesions in both males and females, while the GsdmD^-/-^ mice showed significantly smaller atherosclerotic lesions with ~42% decreased lesion area in females (***Fig. 6C***) and ~33% decreased lesion area in males (***Fig. 6D***), The WT and GsdmD^-/-^ mice injected with control ASO and fed with WTD showed no atherosclerotic plaques (***Fig. S5***).

### GsdmD is cleaved in atherosclerotic plaques and play role in inducing expression of adhesion molecule

To determine if GsdmD is actively cleaved in plaques, we prepared protein extracts from atherosclerotic aortas and performed western blot analysis using the antibody that recognizes only cleaved GsdmD and not the full-length GsdmD. As shown in ***Fig. 7A***, the cleaved GsdmD fragment was detected in the atherosclerotic aortas of WT mice. As expected, the protein extracts of aortas from GsdmD^-/-^ mice showed no positive signal. To investigate the mechanism of GsdmD in promoting atherosclerosis, we also probed the aortic protein extracts for VCAM-1 expression levels. The LDLr ASO injected mice showed higher expression of VCAM-1 in aortic extracts vs. control ASO injected mice (***Fig. 7A***). The aortic extracts from LDLr ASO injected GsdmD^-/-^ mice showed lower induction in levels of VCAM-1 vs. WT mice, indicating that GsdmD expression induces VCAM-1 expression in hyperlipidemic mice. To further confirm the presence of cleaved GsdmD in atherosclerotic plaque areas, we performed immunohistochemistry on aortic sections and stained the tissue sections with cleaved GsdmD antibody and counterstained with hematoxylin. As shown in ***Fig. 7B***, the aortic sections from the LDLr ASO injected WT mice showed a positive signal with cleaved GsdmD antibody while sections from LDLr ASO injected GsdmD^-/-^ mice showed no staining. The plasma ATP levels were shown to be significantly higher in atherosclerotic LDLR^-/-^ mice vs. nonatherosclerotic LDLR^-/-^ mice, and increased plasma ATP exacerbated atherosclerosis in LDLr^-/-^ mice (*31*). The increased plasma ATP levels can serve as a danger signal and induce Nlrp3 inflammasome assembly and IL-1β release in plaques. Increased IL-1β levels can promote increased recruitment of immune cells to plaque area and promote increased ATP release from damaged foam cells due to the release of cytoplasmic content. To determine if the GsdmD pathway contribute to ATP release in atherosclerotic LDLR^-/-^ mice, the plasma ATP levels were determined in WT and GsdmD^-/-^ mice injected with LDLr ASO and fed with atherogenic diet for 9 weeks. As shown in ***Fig. 7C***, the plasma ATP increased significantly in atherosclerotic LDLR^-/-^ mice while the GsdmD^-/-^ LDLR^-/-^ double knockout (KO) mice showed significant attenuation in levels of plasma ATP. These data indicate the decreased IL-1β release in GsdmD^-/-^ LDLR^-/-^ double KO mice can decrease foam cell membrane disintegration and reduce ATP release in plasma.

**Fig. 7.**
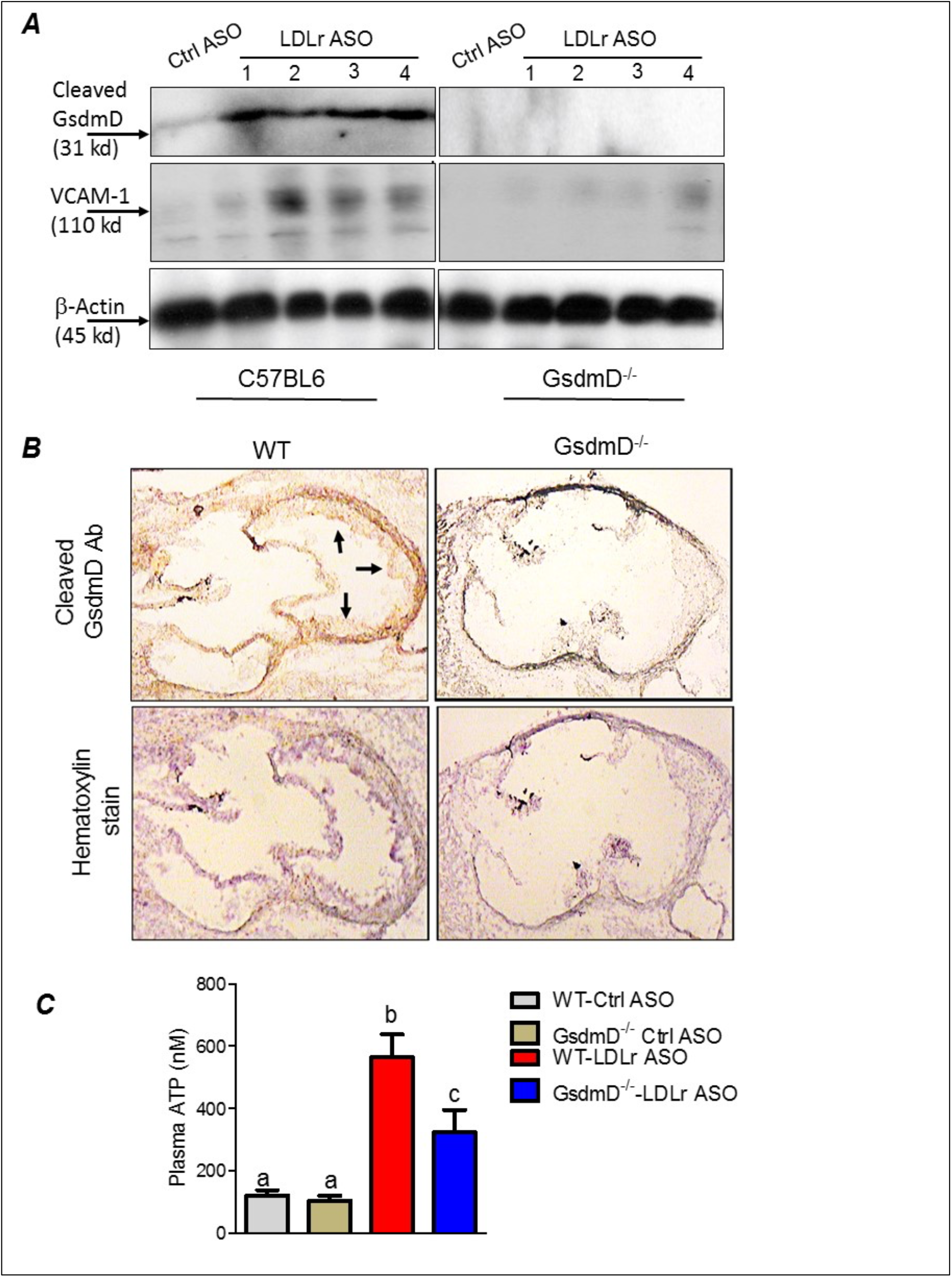
GsdmD is cleaved in atherosclerotic plaques. ***A)*** The 12-weeks old mice were injected with either control or LDLr ASO for 9 weeks and were fed western type diet throughout the course of study. Mice were euthanized after 9-week ASO treatment and protein extracts from aortas of control ASO or LDLr ASO treated mice (N=4) were subjected to western blot analysis using antibody specific for cleaved GsdmD and VCAM-1. GAPDH was probed as a loading control. ***B)*** The fresh Frozen hearts were sectioned, followed by antigen retrieval and endogenous peroxidase activity blocking, and probing with antibody specific for cleaved N-terminal fragment of Gasdermin D. ***C)*** ATP levels in plasma of control ASO or LDLr ASO treated mice fed with atherogenic diet were determined by ATP quantification kit, following manufacturer’s instructions. (mean ± SD, N = 8; different letters show p<0.001, by ANOVA Bonferroni posttest).

## Discussion

Chronic inflammation is a key player in promoting atherosclerosis and cardiovascular disease (*32*). The link between cholesterol efflux by ABC transporters, inflammatory cytokines, and heart disease is becoming clearer with studies showing that inflammation can promote atherosclerosis by dampening the RCT pathway (*15, 33*). Many studies have highlighted the role of Nlrp3 inflammasome/IL-1β pathway in promoting CVD (*1, 3, 4, 25, 34*) and the link between inflammation and CVD is further highlighted by the link between sepsis survivor and higher risk of having a heart attack or stroke (*35*). The role of GsdmD in sterile inflammatory diseases, such as atherosclerosis, is not yet clear. We found that in-vivo Nlrp3 inflammasome assembly in mice by LPS+ATP injections leads to GsdmD -dependent increase in IL-1β levels and reduced reverse cholesterol transport (***Fig. 1, 4***). The macrophages lacking GsdmD were more resistant to inflammation-induced defects in cholesterol efflux (***Fig. 2***). The induction of Nlrp3 inflammasome and ensuing membrane damage due to pore generation by GsdmD can reduce the cell’s ability to efficiently transport cholesterol via the ABCA1-apoA1 pathway. To circumvent the membrane damage contribution towards reduced cholesterol efflux, we used conditioned media isolated from WT BMDMs treated with LPS+ATP to show can reduction in cholesterol efflux. These data pointed toward a more direct role of IL-1β in inhibiting cholesterol efflux. Binding of IL-1β to IL-1 receptor (IL-1r) may activate or block cellular signaling pathways involved in apoA1-ABCA1 cholesterol efflux pathway. The identity of these signaling pathways remains to be elucidated. Another possibility is the disruption of PIP2 trafficking in WT macrophages by the Nlrp3 inflammasome. ABCA1 was identified as the first PIP2 floppase, translocating PIP2 from the inner to outer leaflet of the plasma membrane (*27*). The ABC transporter mediated PIP2 flop was later confirmed by other studies (*28, 36*). We have shown before that treatment of macrophages with FDA approved antileishmanial drugs Miltefosine can cause PIP2 mislocalization from the inner leaflet of plasma membrane (*37*). Miltefosine treated macrophages showed defective Nlrp3 inflammasome assembly and reduced IL-1β release (*37*). Cleaved GsdmD and cleaved IL-1β were reported to bind PIP2 directly, and this binding may retain PIP2 on the inner leaflet and thus reduce PIP2 on the cell surface in the WT macrophages. The GsdmD^-/-^ macrophages on the other hand may retain the ability to translocate PIP2 to the cell surface, and thus sustain more cholesterol efflux activity. In support of role of PIP2 in modulating pyroptosis, the cells expressing ABCA1 or PIP2 binding pleckstrin homology domain showed reduced GsdmD-NT mediated cell lysis (***Fig. 2F, 2G***).

GsdmD^-/-^ mice showed significantly reduced atherosclerosis in LDLr ASO generated hyperlipidemic mice, indicating that GsdmD promotes atherosclerosis in hyperlipidemic mice (***Fig. 6***). Recent studies in apoE^-/-^ KO mice using chemical or adenovirus associated inhibition of GsdmD expression also showed a reduction in atherosclerosis (*38, 39*). GsdmD may affect atherosclerotic plaque progression via several different mechanisms. One of the mechanisms could a simple blockage of IL-1β and IL-18 release from inflamed immune cells. This mechanism is supported by the outcome of CANTOS trial showing that anti-IL-1 β antibodies reduced MACE in CVD patients (*4*) and by a previous study showing the role of IL-18 in atherosclerosis in mice (*40*). Another mechanism by which GsdmD can promote atherosclerosis could be via regulating pyroptosis (*18–20*). Unlike apoptosis, which has been a main focus of studies aimed at understanding macrophage cell death in plaque area due to cholesterol overload, the role of pyroptotic cell death in atherosclerosis is not clear. The presence of cleaved N-terminal fragment of GsdmD in aortic tissue from atherosclerotic plaque-bearing mice may indicate the role of pyroptotic cell death in the progression of atherosclerosis, but is not a conclusive evidence as GsdmD can also get cleaved in living macrophages (*7*). The clearance of apoptotic cells by phagocytosis in plaque is of high importance as defective efferocytosis promotes further inflammation and plaque progression (*41*). The pyroptotic cells may not be cleared by efferocytosis as effectively as apoptotic cells due to the lack of proper “eat-me signal”, which is an exposure of phosphatidylserine (PS) on the cell surface. Importantly, a recent study has shown the existence of a robust apoptotic caspase network that is activated in parallel to GsdmD-mediated plasma membrane permeabilization and shifts the balance between apoptotic and pyroptotic macrophage cell death (*42*). We found that lipid-laden GsdmD KO foam cells exhibit higher apoptotic cell death vs. WT macrophages (***Fig. 3C***). Thus, KO of GsdmD may reduce atherogenic pyroptotic cell death, and may tilt the balance toward athero-protective apoptotic cell death.

Neutrophil extracellular traps (NETs) play important role in trapping extracellular pathogens but they can promote atherosclerosis by activating macrophages to release more pro-inflammatory cytokines. Hyperlipidemia and cholesterol crystals can promote the formation of NETs that in turn can play role in the progression of atherosclerosis (*43, 44*). Interestingly, GsdmD is also involved in NETosis and is proteolytically activated by Neutrophil proteases (*5*). Thus, GsdmD KO may dampen the progression of atherosclerosis due to reduced formation of NETs. Further studies are required to decipher if the GsdmD atherogenic activity is mediated solely by macrophages or other immune cells such as neutrophils, B-cells, or T-cells, also contribute to GsdmD atherogenic activity. Recently, AIM2 inflammasome, caspase 1 and 11, and GsdmD were shown to play in role in clonal haematopoiesis mediated exacerbation of atherosclerosis(*45*). Clonal haematopoiesis increases the risk of myocardial infarction and stroke independently of traditional risk factors(*46, 47*). The JAK2V617F (JAK2VF) mutation leads to clonal haematopoiesis via increased JAK–STAT signaling, Using mice that express Jak2VF selectively in macrophages, Gsdmd deletion was shown to reduce necrotic core formation and increase cap thickness, but showed no change in overall lesion area (*45*). Similarly, the bone-marrow transplanted Gsdmd^-/-^ into Ldlr^-/-^ control mice did not change the lesion area (*45*). These data and our study indicate that GsdmD may also have a non-myeloid role in promoting atherosclerosis. Previous studies have shown the major role of vascular smooth muscle cells (VSMCs) in promoting atherosclerosis, with data showing that VSMCs can adopt phenotypes resembling foam cells and macrophages(*48–50*). The whole-body knockout of GsdmD may alter the response of VSMCs to lipids and other inflammatory stimuli to modulate progression of atherosclerosis. Similarly, the GsdmD KO may affect adaptive responses of endothelial cells, including but not limited to autophagy and pyroptosis, to lipid-burden and cytokines released from inflamed foam cells.

Our data show a novel role of GsdmD in the progression of atherosclerosis, independent of lipid levels. The idea of combining lipid-lowering therapeutics with anti-inflammatory drugs is getting increasingly recognized (*51*), and targeting GsdmD in combination of LDL lowering therapies such as statins or PCSK9 antibody may serve as potential therapeutic to treat atherosclerosis and cardiovascular disease.

### Highlights of this work

The Nlrp3/AIM2 inflammasome target protein “GsdmD” is involved in multiple pro-inflammatory pathways such as the release of IL-1β, pyroptotic cell death, and NETosis. Our data provide the first evidence for the role of GsdmD in promoting atherosclerosis. Thus, targeting GsdmD in conjunction with current LDL lowering therapies may provide additional benefits compared to anti-IL-1β therapy, via mechanisms of decreasing pyroptosis and NETosis.

- Macrophages lacking GsdmD are more resistant to defects in cholesterol efflux after inflammasome activation.
- Treatment of macrophages with conditioned media isolated from inflamed WT, but not the GsdmD^-/-^ macrophages, decreased apoA1 binding to the cells.
- ABCA1 expression reduced the binding of N-terminal fragment of GsdmD to the cell membrane and prevented cell lysis. These effects are attributed to ABCA1’s PIP2 floppase activity.
- The conditioned media from inflammasome-activated WT, but not the GsdmD^-/-^ macrophages, promotes foam cell formation.
- Inflammasome-activated GsdmD^-/-^ macrophages are more susceptible to apoptotic cell death.
- GsdmD^-/-^ mice show blunted suppression of RCT in vivo (plasma, liver, feces) upon inflammasome activation.
- GsdmD is cleaved in aortic root plaques and LDLR ASO treated GsdmD^-/-^ mice show reduced VCAM1 expression in aortic root plaques and decreased atherosclerotic lesions.

## Material and Methods

### Mice and diets

All animal experiments were performed in accordance with protocols approved by the Cleveland State University and Cleveland Clinic Institutional Animal Care and Use Committee. The C57BL6J mice were purchased from the Jackson Laboratory and the GsdmD^-/-^ KO-C57BL/6J mice were generated earlier (*52*) and kindly provided by Dr. Russell Vance (UC, Berkeley). The GsdmD^-/-^ KO genotype was confirmed by PCR & sequencing of Exon 2 of GsdmD using primers, GsdmD Fwd: ATAGAACCC GTGGAGTCCCA and GsdmD Rev: GGCTTCCCTCATTCAGTGCT as described earlier (*52*). The 12-week-old mice, males and females, in C57BL/6J or GsdmD^-/-^ - C57BL/6J backgrounds were maintained in a temperature-controlled facility with a 12-h light/dark cycle. Mice were given free access to food and water. The standard chow diet (SD, 20% kcal protein, 70% kcal carbohydrate and 10% kcal fat, Harlan Teklad) was used for regular maintenance and breeding. For generating hyperlipidemia for atherosclerosis studies, mice were fed an atherogenic Western type diet (WD) (ENVIGO, 0.2% cholesterol with 42% adjusted calories from fat, TD.88137).

### ASO injections

GalNAc-conjugated Gen 2.5 ASO targeting mouse Low-density Lipoprotein receptor (LDLr) and control ASO were kindly provided by Adam Mullick from Ionis Pharmaceuticals. The. ASO treatment started at week 12 with i.p. injection of 5 mg/Kg body weight, 1 x per week until sacrifice at week 21. The oligo stock was prepared at 500mg/ml in sterile saline, such that a 25 g mouse would receive a 250 μl injection. The control group receive 1x per week control ASO. Following the ASO injection for 9 weeks, animals were sacrificed and hearts were perfused for sectioning for quantification of atherosclerotic lesions.

### Isolation of Bone marrow derived macrophages

WT C57BL/6 or C57BL/6 *GsdmD KO* mice were maintained on standard chow diet and water. Mice were euthanized by CO2 inhalation and femoral bones were removed and flushed for marrow for isolating bone-marrow cells Detailed method for preparing bone-marrow derived macrophages is described in *Supplementary material and methods section.*

### In-vivo Nlrp3 inflammasome assembly and IL-1β release assays

The 12-week-old WT or GsdmD^-/-^ mice were IP injected with either 5 μg LPS, or sterile PBS. After 4h of LPS or PBS injection, the Nlrp3 inflammasome assembly in mice was induced by IP injection of ATP (0.5 ml of 30 mM, pH 7.0). The mice were euthanized after 30 min and peritoneal cavity was lavaged with 5 ml PBS. Approximately 3.5 ml peritoneal lavage fluid was recovered from each mouse and centrifuged at 15K for 10 min at room temperature. The supernatant was subjected to IL-1β ELISA, using mouse IL-1b Quantikine ELISA kit (MLB00C, R & D systems) and following manufacturer’s instructions.

### Cholesterol efflux assay

Cholesterol efflux assays were performed as described earlier (*27*). The WT or GsdmD^-/-^ BMDMs were radiolabeled with 0.5 μCi/ml of [3H]-cholesterol, ABCA1 expression in BMDMs was induced by treatment with T0901317, and chase was performed with serum free media containing cholesterol acceptor apoA1. The percent cholesterol efflux was calculated as 100 × (medium dpm) / (medium dpm + cell dpm). Detailed method is described in *Supplementary material and methods section.*

#### Mice RCT assay

WT Murine bone marrow derived macrophages were incubated with tritium labeled [3H] cholesterol and acetylated LDL to generate foam cells. The day prior to radiolabeled foam cell injection, mice received intraperitoneal injections of PBS or 5μg/ml LPS and three hours later mice treated with LPS received 30 mM of adenosine tri-phosphate, (sigma) pH=7.5-8. Mice received daily LPS/ATP injection through the course of the study. RCT to the plasma, liver, and feces was calculated as the % (dpm appearing in plasma/total dpm injected). Detailed method is described in *Supplementary material and methods section.*

### Western blotting

BMDMs or RAW264.7 cells were grown and treated as indicated. The PBS-washed cell pellet was lysed in MPER or RIPA lysis buffer supplemented with protease inhibitors and PMSF. Detailed method is described in *Supplementary material and methods section.*

### Cholesterol measurements

Total cholesterol was measured by using Stan Bio Total cholesterol reagent (#1010-225), following manufacturer’s instructions.

### Atherosclerotic lesion quantification

Mice were sacrificed by CO2 inhalation and weighed at 21 weeks of age. Whole blood was collected from the retroorbital plexus into a heparinized glass capillary, mixed with EDTA and spun in a microfuge to obtain plasma. The circulatory system was perfused with 10 mL PBS and the heart was excised and preserved in 10% phosphate buffered formalin. The quantitative assessment of atherosclerotic plaque area in the aortic root was performed as previously described (*53*). Lesion areas were quantified as the mean value in multiple sections at 80 μm intervals using Image Pro software (Media Cybernetics).

### Statistics

All statistics were performed using GraphPad Prism software. Comparison of two groups was performed by two-tailed *t*-test and comparison of more than two groups was performed by ANOVA with the specified posttest. Values shown are mean ± SD.

## Non-standard abbreviations and acronyms

AcLDL: acetylated-low-density lipoprotein
AIM2: absent in melanoma 2
ASO: antisense oligonucleotides
CAD: coronary artery disease
CANTOS: canakinumab anti-inflammatory thrombosis outcomes study
CVD: cardiovascular disease
GsdmD: gasdermin D
IL-1β: interleukin 1-beta
IL-18: interleukin 18
LDLr: low-density lipoprotein receptor
MACE: major adverse coronary events
NLRP3: nod-like receptor family pyrin domain-containing 3
NETs: neutrophil extracellular traps
(oxLDL: oxidized low-density lipoprotein
PIP2: phosphatidylinositol 4,5-bisphosphate
PS: phosphatidylserine
TLR: toll-like receptorx

## Author Contributions

K.G. designed and directed research. E.O, C.A.T, D.Z, A.J.I, J.H, M.K. and K.G performed research. E.O, C.A.T, and K.G. analyzed data. K.G. drafted the manuscript. All authors critically reviewed the manuscript.

## Acknowledgments

This work was supported by National Institutes of Health Grant RO1HL148158 (to K.G), American Heart Association Scientist Development Grant SDG25710128 (to K.G), and Cleveland State University startup funds (to K.G.). C.A.T is supported by F31HL134231 and J.D.S is supported by RO1HL128268 and R01 HL130085.

