## Supplementary for "Gasdermin D mediates inflammation-induced defects in reverse cholesterol transport and promotes atherosclerosis"

Running Title: GsdmD mediates inflammation induced defects in RCT and promotes atherosclerosis.

Emmanuel Opoku[1,4], C. Alicia Traughber[1,2,3,4], David Zhang[1], Amanda J Iacano[1], Mariam Khan[2,3], Juying Han[1], Jonathan D Smith[1,] and Kailash Gulshan*[1,2,3]

### *SI Materials and Methods.*

#### *Material:*

#### *Table S1: Antibodies & reagents*

#### *Antibodies*

| | | |
|---|---|---|
| Novus Bio | ABCA1 | NB400-05 |
| Cell signaling: | GAPDH (14C10) | 2118 |
| Cell signaling | NLRP3 (D4D8T) | 15101 |
| Cell signaling | Cleaved GSDMD (Asp276) | 50928 |
| Cell signaling | IL-1β | 12507 |
| Cell signaling | Beta-Actin | 8457 |
| Cell signaling | VCAM-1 | 32653 |
| R & D systems/<br>Santa Cruz Biote | LDLr | AF2255/<br>SC 18823 |
| BioRad | Goat Anti-Rabbit IgG (H + L)-HRP Conjugate | 1706515 |

#### *Reagents*

| | |
|---|---|
| Western Type Diet (42% Calories from Fat) | Teklad |

| | |
|---|---|
| LDLR anti-sense oligonucleotides (ASO) | Ionis Pharmaceuticals |
| heparinize micro-hematocrit capillary tubes | Fisher #22-362-566 |
| O.C.T | Fisher # 23-730-571 |
| $H_2O_2$ | Sigma #H1009 |
| PAP pen for immunostaining (5mm tip width) | Millipore Sigma #Z377821 |
| Normal Goat Serum | Vector Labs #S-1000 |
| R & D systems | Mouse rIL-1b#401-ML-025 |
| Triton X-100 | ICN Biomedicals |
| Vector® NovaRED® Substrate Kit, Peroxidase (HRP) | Vector Labs #SK-4800 |
| Hematoxylin Solution, Harris Modified | Sigma #HHS32 |
| Aqua-Mount® Mounting Medium | Fisher # 41799-008 |
| Thermo Scientific™ Richard-Allan Scientific™ Cover Glass | Fisher #22-050-222 |
| 26G and 27G needles | BD Biosciences |

***Methods***

**Western blotting:** BMDMs were grown and treated as indicated. The PBS-washed cell pellet was lysed in MPER lysis buffer supplemented with protease inhibitors. After discarding the nuclear pellet, the protein concentration was determined using the BCA protein assay (Pierce). 10-50 µg of cell protein samples were resolved on Novex 4-20% Tris-Glycine Gels (Invitrogen) and transferred onto polyvinylidene fluoride membranes (Invitrogen). Blots were incubated sequentially with 1:1000 rabbit polyclonal antibody raised against Nlrp3 (cell signaling), or 1:1000 rabbit polyclonal antibody raised against

IL-1β (cell signaling) and 1:15,000 horseradish peroxidase-conjugated goat anti-mouse secondary antibody (Biorad) were used. The signal was detected with an enhanced chemiluminescent substrate (Pierce). The beta-actin was used as a loading control.

**Isolation of Bone marrow derived macrophages:** WT C57BL/6 or C57BL/6 *GsdmD* $^{-/-}$ *KO* mice were maintained on standard chow diet and water. Mice were euthanized by $CO_2$ inhalation and femoral bones were removed. The marrow was flushed out of the bones into a 50 ml sterile tube using a 10 ml syringe with a 26-gauge needle filled with sterile DMEM. Cells were centrifuged for 5 min at 1,800 rpm at 4°C, followed by two washes with sterile PBS. The cells were resuspended in sterile-filtered BMDM growth media (DMEM with 7.6% fetal bovine serum, 15% L-cell conditioned media, and 0.76% penicillin/streptomycin mixture) and plated in 10 cm culture dishes and incubated at 37°C for 14 days. Cell media was replaced every 2–3 days for 2 weeks. The cells were routinely visualized under microscope for proliferation and differentiation into confluent BMDMs.

**Cholesterol efflux assay**: The BMDMs were plated in 24-well plates at a density of 300-400,000 cells per well. The cells were labeled with 0.5 µCi/ml of [3H]-cholesterol (PerkinElmer, Waltham, MA) in DMEM containing 1% FBS for 24h. The labeled cells were induced for ABCA1 expression for 16h with 0.5 µM T0901317 (Sigma). The labeled cells were induced for Nlrp3 inflammasome assembly by LPS/Nigericin treatment or left untreated as control. The cells were washed twice with serum free media and chased with cholesterol acceptor 5μg/ml apolipoprotein A1 (apoA1) for 6h. The chase media was collected and subjected to brief centrifugation to pellet any residual debris. The cleared media was used to determine the radioactive counts effluxed out of the cells. Radioactivity in the cells was determined by extraction in hexane:isopropanol (3:2) with the solvent evaporated in a scintillation vial prior to counting. The percent cholesterol efflux was calculated as 100 × (medium dpm) / (medium dpm + cell dpm).

***Mice RCT assay***. WT murine bone marrow macrophages were cultured for 11 to 14 days in DMEM supplemented with 20% L-cell conditioned medium, 10% fetal bovine serum and 1% penicillin/streptomycin. To generate foam cells [3H] cholesterol ((2 μCi/ml; Perkin Elmer, Norwalk, USA) was incubated with 100 μg/ml acetylated LDL, then combined with DMEM containing 20% L-cell conditioned medium) to load and label murine bone marrow macrophages or 24 hours. The day prior to radiolabeled foam cell injection, mice received intraperitoneal injections of PBS or 5μg/ml LPS and  Three hours later mice treated with LPS received 30 mM of adenosine tri-phosphate, (sigma) pH=7.5-8.  Mice received LPS/ATP through the course of the study. They day of foam cell injection, cells were washed twice with DMEM prior to harvesting for in vivo injection, and ~2 million [3H]cholesterol dpm in a volume of 0.25 ml were injected subcutaneously on the upper back of recipient mice.  At 24, 48, and 72 hours post cell injections, plasma and feces were collected from mice.  Plasma radioactivity was determined, and total plasma dpm was calculated by estimating blood volume to be equal to 7% of the body weight and plasma to be 55% of the blood volume. RCT to the plasma was calculated as the % (dpm appearing in plasma/total dpm injected). Collected feces were allowed to dry overnight at 55°C and then weighed. Feces were then hydrated in 10 ml of 50% ethanol solution followed by homogenization then an internal recovery standard of 10,000 dpm of [14C]cholesterol (Perkin Elmer, Norwalk,

USA) was added to each sample. The radioactivity was quantified as described in detail above. Upon sacrifice at 72h, the liver was removed and weighed. A piece of liver, ~0.2 g, was isolated, weighed, suspended in PBS, homogenized, and a known amount of [14C]cholesterol radioactivity was added as a recovery standard. The radioactivity in an aliquot of 0.3 ml of the liver homogenate was measured by liquid scintillation counting The [14C]cholesterol dpm was used to back calculate the [3H] recovery for the entire liver homogenate, which was further used to calculate the total amount of radioactivity in the liver. In addition to [14C] cholesterol internal standard, all RCT data was standardized to mouse body or fecal weight each harvest time point. Outliers in the RCT study usually attributed to leakage of injected cell volume were excluded from analysis.

**Immunohistochemistry**

Fresh Frozen hearts were embedded in O.C.T (Fisher # 23-730-571). Ten micrometer sections of the aortic root were cast onto superfrost microscope slides (Fisher # 12-550-15) and stored at -80°C. For immunostaining, frozen tissue slides were placed at room temperature for 2 minutes then fixed with ice-cold 100% acetone, followed by 3 PBS washes. Next, endogenous peroxidase activity was blocked with 3% $H_2O_2$ (Sigma #H1009) prepared in methanol, followed by 3 PBS washes. Tissue sections were outline using a hydrophobic marker (Millipore Sigma #Z377821) and then blocked for 1h with 5% goat serum (Vector Labs #S-1000) containing 0.3% Triton X-100 (ICN Biomedicals) in PBS. Sections were then probed with a 1:50 dilution of cleaved Gasdermin D antibody (Cell Signaling Technology #50928) at 4°C overnight, then washed 3 times with PBS. Tissues were then incubated with 1:1000 a goat anti-rabbit secondary (BioRad #1706515) for 30 minutes, then washed with PBS 3 times. Staining was developed by the Vector® NovaRED® Substrate Kit, Peroxidase (HRP) (Vector Labs #SK-4800) according to manufacturer instructions. Sections were then washed with water and counterstained with hematoxylin (Sigma #HHS32). Section were mounted with mounting media (Fisher # 41799-008), covered by glass coverslip (Fisher #22-050-222). Once dry, images were captured under light microscopy.

**Supplementary Figure Legends**

***Fig. S1: Effect of rIL-1$\beta$ on cholesterol efflux and controls for Fig. 2B***. **A)** BMDMs isolated from WT or GsdmD$^{-/-}$ mice were loaded with $^3$H-cholesterol + 50$\mu$g/ml AcLDL for 24h and treated with LXR-agonist T0901317 (Sigma-Aldrich) to induce ABCA1 expression. The Nlrp3 inflammasome was induced by LPS+ Nigericin treatment or cells were treated with LPS + 10 ng/ml recombinant mouse IL-1$\beta$ (R & D systems), followed by cholesterol efflux assay using lipid-free apoA1 (5$\mu$g/ml) in serum-free DMEM as chase media for 4h at 37$^o$C. Values are % cholesterol efflux (mean ± SD, N = 4, different letters represent p<0.01 by ANOVA Bonferroni posttest). **B)** *Controls for Fig 2B*; controls without ABCA1 expression (not treated with T-compound) are presented along with samples presented in Fig. 2B. The BMDMs isolated from WT or GsdmD$^{-/-}$ mice were loaded with $^3$H-cholesterol + 50$\mu$g/ml AcLDL for 24h. The BMDMs were treated with conditioned media isolated from LPS/ATP treated WT or GsdmD$^{-/-}$ BMDMs ± LXR-agonist T0901317 for 24h. The cholesterol efflux assay was performed using lipid-free apoA1 (5$\mu$g/ml) in serum-free DMEM as chase media for 4h at 37$^o$C. Values are % cholesterol efflux. C**)** BMDMs isolated from WT mice were loaded with $^3$H-cholesterol + 50$\mu$g/ml AcLDL for 24h and treated with LXR-agonist T0901317 (Sigma-Aldrich) to induce ABCA1 expression. The cells were treated with LPS + different doses of recombinant mouse IL-1$\beta$ (R & D systems), followed by cholesterol efflux assay using lipid-free apoA1 (5$\mu$g/ml) in serum-free DMEM as chase media for 4h at 37$^o$C. Values are % cholesterol efflux (mean ± SD, N = 4. **** represent p<0.0001, ** represent p<0.01 by ANOVA Bonferroni posttest

***Fig. S2: ABCA1 expression in BMDMs± conditioned media:*** Western blot of ABCA1 was performed for cell extracts of T-compound treated WT or GsdmD$^{-/-}$ BMDMs treated with conditioned media from either WT control BMDMs, WT BMDMs +LPS+ATP, or GsdmD$^{-/-}$ BMDMs + LPS+ATP. The primary antibody used was Rabbit anti-ABCA1 (1:500 dilution) and secondary antibody was goat anti-rabbit (1: 10,000 dilution), and GAPDH was used as a loading control.

***Fig. S3: Localization of PIP2 binding PH-PLC domain in HEK293 cells stably transfected with 2X-PH-PLC-eGFP construct:*** Plasma membrane localization of eGFP tagged PIP2 binding pleckstrin homology (PH) domain of phospholipase- C

(PLC). The HEK293 cells stably transfected with 2X-PH-PLC-eGFP construct are imaged by fluorescent microscopy.

***Fig. S4: Original micrograph of figure 3A***. WT and GsdmD$^{-/-}$ BMDMs were treated with 25ug/ml oxLDL in non-conditioned, WT BMDM conditioned, or GsdmD$^{-/-}$ conditioned media for 24h, then stained with Nile red. Foam cell formation was assessed by microscopy and quantified by flow cytometry. Conditioned media generated by treating BMDMs with 1ug/ml LPS for 3h and 5mM ATP for 30m.

***Fig. S5: Control ASO did not generate hyperlipidemia or atherosclerotic lesions in WTD fed WT or GsdmD$^{-/-}$ KO mice***. The 12-week old mice from both sexes were injected with either control ASO for 9 weeks. ***A***) Total plasma cholesterol levels were determined for both WT-week 0 vs. WT-week 9 and GsdmD$^{-/-}$-week 0 vs. GsdmD$^{-/-}$-week 9, N=5, mean $\pm$ S.D, n.s= non-significant by two-tailed t-test).
Representative images from showing aortic root lesions from WT or GsdmD$^{-/-}$ KO mice injected with control ASO or LDLr ASO and fed WTD, stained with Oil red O and hematoxylin.

*Figure S1*

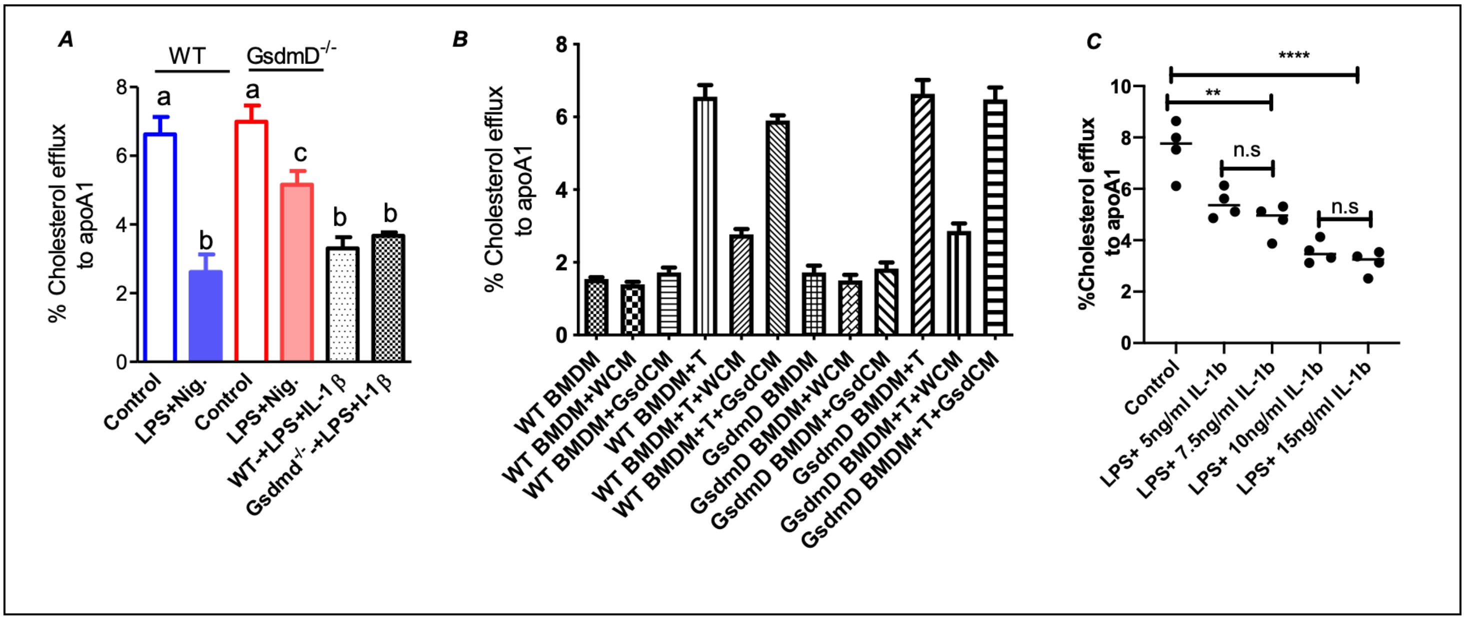
A
WT
GsdmD-/-
% Cholesterol efflux to apoA1
a
b
a
c
b
b
Control
LPS+Nig.
Control
LPS+Nig.
WT-+LPS+IL-1 β
Gsdmd-/- -+LPS+I-1 β
B
% Cholesterol efflux to apoA1
WT BMDM
WT BMDM+WCM
WT BMDM+GsdCM
WT BMDM+T
WT BMDM+T+WCM
WT BMDM+T+GsdCM
GsdmD BMDM
GsdmD BMDM+WCM
GsdmD BMDM+GsdCM
GsdmD BMDM+T
GsdmD BMDM+T+WCM
GsdmD BMDM+T+GsdCM
C
%Cholesterol efflux to apoA1
****
**
n.s
n.s
Control
LPS+ 5ng/ml IL-1b
LPS+ 7.5ng/ml IL-1b
LPS+ 10ng/ml IL-1b
LPS+ 15ng/ml IL-1b

***Figure S2***

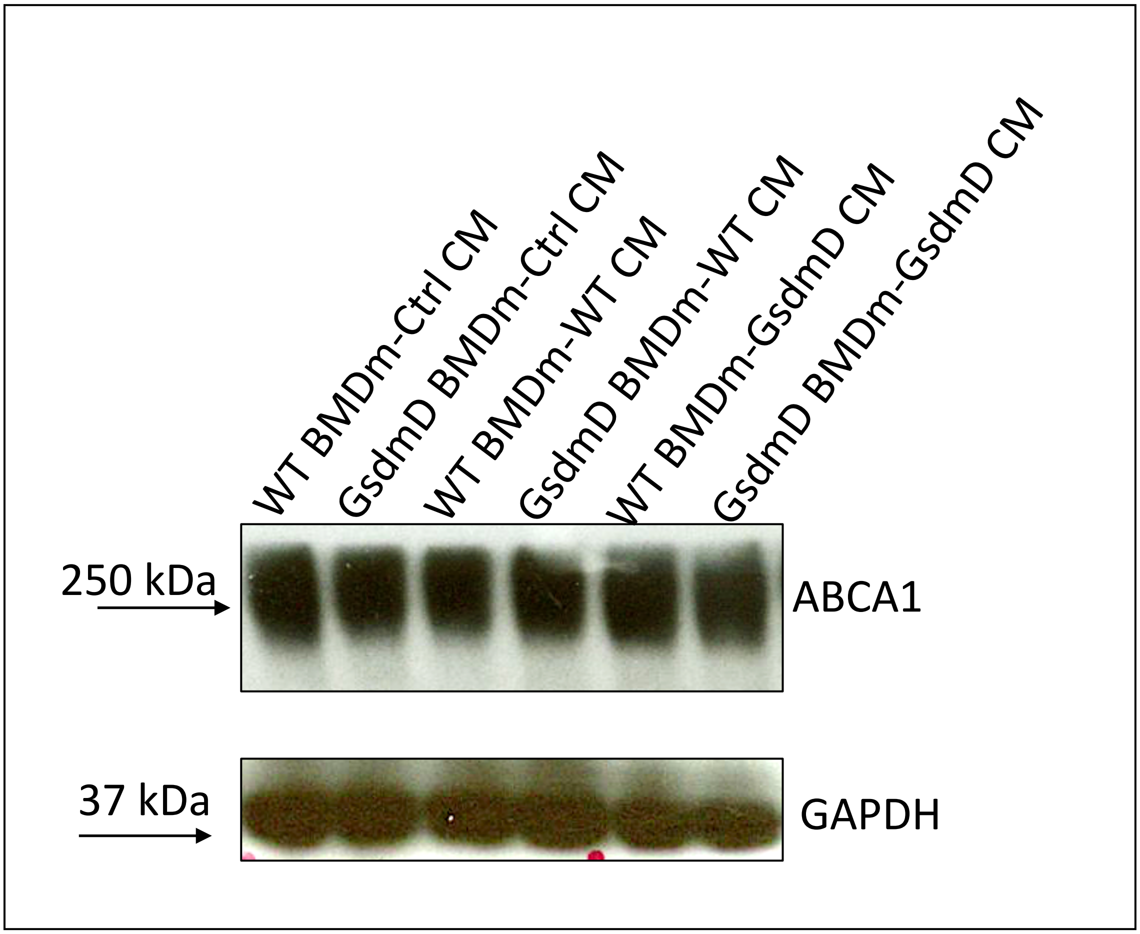
WT BMDm-Ctrl CM
GsdmD BMDm-Ctrl CM
WT BMDm-WT CM
GsdmD BMDm-WT CM
WT BMDm-GsdmD CM
GsdmD BMDm-GsdmD CM
250 kDa
ABCA1
37 kDa
GAPDH

*Figure S3*

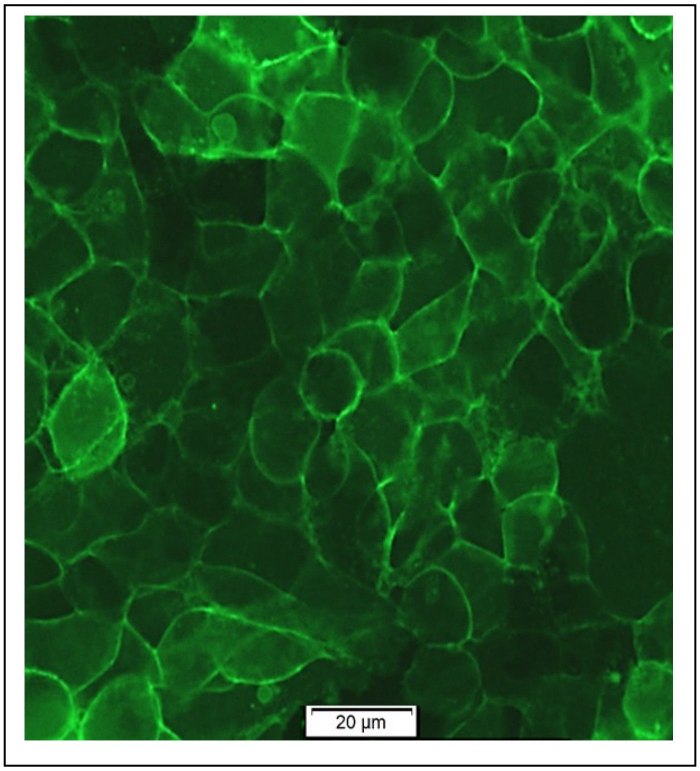

*Fig. S4*

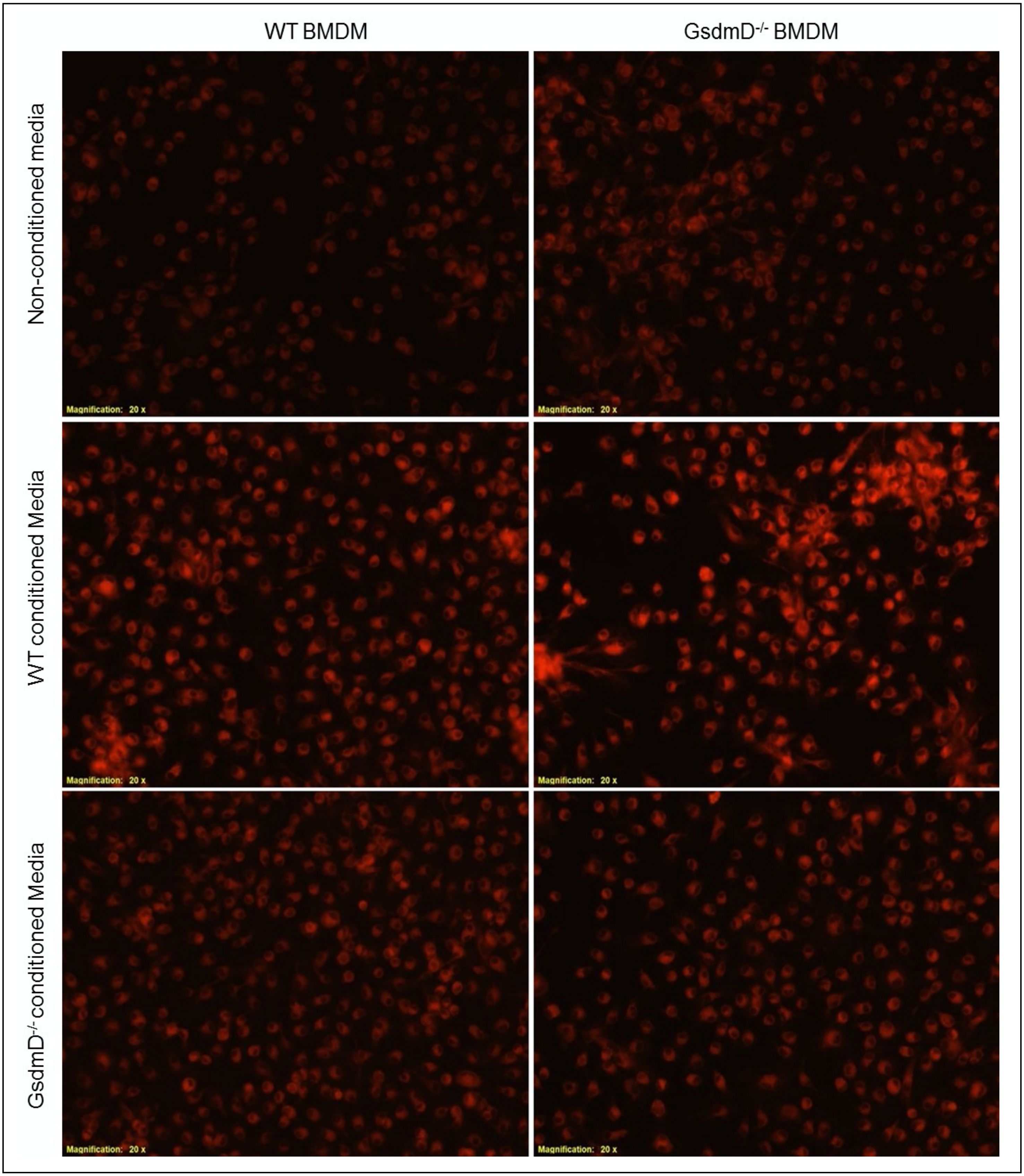
WT BMDM
GsdmD-/- BMDM
Non-conditioned media
WT conditioned Media
GsdmD-/- conditioned Media
Magnification: 20 x
Magnification: 20 x
Magnification: 20 x
Magnification: 20 x
Magnification: 20 x
Magnification: 20 x

***Figure S5***

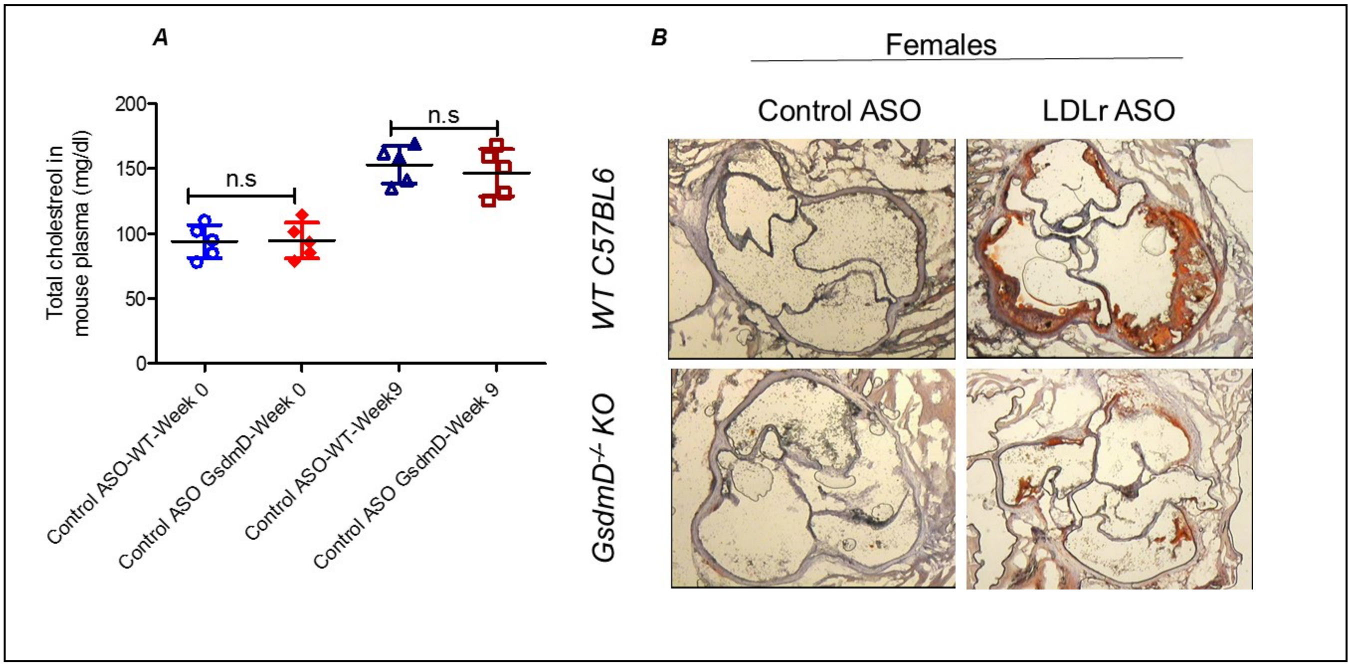
A
B
Total cholestreol in mouse plasma (mg/dl)
200
150
100
50
0
n.s
n.s
Control ASO-WT-Week 0
Control ASO GsdmD-Week 0
Control ASO-WT-Week9
Control ASO GsdmD-Week 9
Females
Control ASO
LDLr ASO
WT C57BL6
GsdmD-/- KO
